# *Osterix* functions downstream of anti-Müllerian hormone signaling to regulate Müllerian duct regression

**DOI:** 10.1101/237529

**Authors:** Rachel D. Mullen, Ying Wang, Bin Liu, Emma L. Moore, Richard R. Behringer

**Affiliations:** Department of Genetics, University of Texas MD Anderson Cancer Center, Houston, Texas 77030

## Abstract

In mammals, the developing reproductive tract primordium of male and female fetuses consists of the Wolffian duct and the Müllerian duct (MD), two epithelial tube pairs surrounded by mesenchyme. During male development, mesenchyme-epithelia interactions mediate MD regression to prevent its development into a uterus, oviduct and upper vagina. It is well established that transforming growth factor-beta family member anti-Müllerian hormone (AMH) secreted from the fetal testis and its type 1 and 2 receptors expressed in MD mesenchyme regulate MD regression. However, little is known about the molecular network regulating downstream actions of AMH signaling. To identify potential AMH-induced genes and regulatory networks controlling MD regression in a global non-biased manner, we examined transcriptome differences in MD mesenchyme between males (AMH signaling on) and females (AMH signaling off) by RNA-Seq analysis of purified fetal MD mesenchymal cells. This analysis found 82 genes up-regulated in males during MD regression and identified *Osterix (Osx)/Sp7*, a key transcriptional regulator of osteoblast differentiation and bone formation, as a novel downstream effector of AMH signaling during MD regression. Osx/OSX was expressed in a male-specific pattern in MD mesenchyme during MD regression. OSX expression was lost in mutant males without AMH signaling. In addition, transgenic mice ectopically expressing human AMH in females induced a male pattern of *Osx* expression. Together these results indicate that AMH signaling is necessary and sufficient for *Osx* expression in the MD mesenchyme. In addition, MD regression was delayed in *Osx* null males, identifying *Osx* as a new factor that regulates MD regression.

**Significance:** In mammals, each embryo forms both male and female reproductive tract organ progenitor tissues. Anti-Müllerian hormone (AMH) secreted by fetal testes acts on mesenchyme cells adjacent to the Müllerian duct (MD) epithelium, the progenitor tissue of the female reproductive tract, to induce MD regression. While AMH and early AMH signaling components are elucidated, downstream gene networks directing this process are largely unknown. A global non-biased approach using whole transcriptome sequencing of fetal MD mesenchymal cells identified 82 factors as potential target genes of AMH including *Osterix (Osx)*. Our findings provide *in vivo* evidence *Osx* is an AMH-induced gene that regulates MD regression. Identification of *Osx* may provide key insights into gene regulatory networks underlying MD regression and male sex differentiation.

## Introduction

In mammals, the Wolffian ducts (WD) differentiate into the male epididymides, vas deferens and seminal vesicles, whereas the Müllerian ducts (MD) develop into the female oviducts, uterus and upper vagina. Reproductive tract differentiation in amniotes is unique because initially both WD and MD are generated in the embryo independent of genetic sex. Sex-specific signaling results in loss of the WD in females and regression of the MD in males (1, 2). MD regression requires transforming growth factor-β family member AMH secreted from the Sertoli cells of the fetal testis and its type 1 and 2 receptors present in MD mesenchyme (3-7). Following AMH binding, AMH type 2 receptor (AMHR2) recruits a type 1 receptor into a heteromeric complex. Within the complex AMHR2 trans-phosphorylates and activates the type 1 receptor kinase. This activation results in the phosphorylation of an R-Smad and formation of an R-SMAD/SMAD-4 complex that translocates into the nucleus to transcriptionally activate AMH signaling pathway target genes. AMH type 1 receptors AVCR2 and BMPR1A, and AMH R-Smad effectors (SMAD1, SMAD5 and SMAD8) function redundantly for MD regression and are shared with the bone morphogenetic protein (BMP) pathway (8, 9).

While the upstream components (AMH, Type 1 & 2 receptors and R-Smads) of AMH signaling are well known, the downstream transcriptional effectors of AMH signaling are still largely unidentified. To date, only the WNT pathway effector *β-catenin (Ctnnb1)* has been shown to be required for AMH-induced MD regression *in vivo* (10). The requirement of *Ctnnb1* for MD regression suggests WNT signaling is important for the downstream actions of AMH during MD regression in males. However candidate gene approaches have failed to identify an individual WNT, WNT effector or WNT regulator whose *in vivo* loss blocks MD regression (10-12).

Because of the limited success of candidate gene approaches in uncovering AMH signaling effectors, in the current study we undertook a non-biased global approach using next generation transcriptome sequencing. To elucidate potential gene networks and novel AMH signaling targets, we used RNA-Seq analysis of YFP (yellow fluorescent protein)-positive MD mesenchymal cells flow sorted from embryonic day (E) 14.5 *Amhr2*^*Cre/*+^*; R26R*^*yfp*/+^ reproductive tracts to identify transcriptome differences between males and females during regression. This analysis identified *Osterix (Osx)/Sp7* as a downstream effector of AMH signaling during MD regression. Our results indicate that AMH signaling is necessary and sufficient for *Osx* expression in the MD mesenchyme and *Osx* regulates the timing of MD regression.

## Results

### Transcriptome analysis identifies candidate genes upregulated in male MD mesenchyme during regression

During embryogenesis, *Amhr2* is expressed in the MD mesenchyme of male and female fetuses, but *Amh* is only expressed in males. Thus, the AMH signaling pathway is only active in male fetuses. To isolate male and female MD mesenchymal cells, *Amhr2-Cre* mice, which express Cre recombinase in the MD mesenchyme and somatic cells of the gonad, were bred to *R26R-YFP* reporter mice. E14.5 was chosen for this analysis because AMH has been expressed for 2 days in males, there are changes in the MD mesenchyme and MD epithelium, but no breaks or gaps in the MD have occurred. Mice carrying the Cre and YFP alleles were identified by fluorescence and sex was determined by gross morphology of the gonad using a dissecting microscope. The mesonephros/gonad complex was isolated and the gonads removed. Two or more mesonephroi were pooled and digested in trypsin and the resulting single cell suspensions were then sorted for YFP+ cells using FACS (**Fig. 1A-C**). cDNA libraries were generated from three biological replicates each for male and female. Indexed libraries were pooled and sequenced on the Illumina HiSeq2000 platform (**Fig. 1D**). ~20 million paired end 76 base pair reads were obtained for each library, of which ~70% mapped to at least one location in the mouse reference genome build NCBIM37. Mapped mouse RNA-seq data were then subjected to differential expression analysis, using Cufflinks (13) and DESeq (14). Genes with significant differential expression were then further parsed by removing genes with fragments per kilobase of exon per million reads mapped (FPKM) at or below those for *Amh* in males, a gene known to be expressed at low levels in MD mesenchyme (15). These analyses revealed 82 genes that were significantly upregulated in males versus females during MD regression (**Table 1**). IPA analysis predicted both BMP2 and BMP4 as upstream activators (data not shown). This is consistent with activation by the AMH signaling pathway because it shares components of the BMP signaling pathway, including type 1 receptors and R-SMADs. In the IPA analysis, *Osx* was in our gene list, predicting BMP as an upstream activator for our dataset. This suggested *Osx* as a potential downstream effector of AMH signaling and led us to analyze *Osx* expression and function during MD regression.

**Fig. 1.**
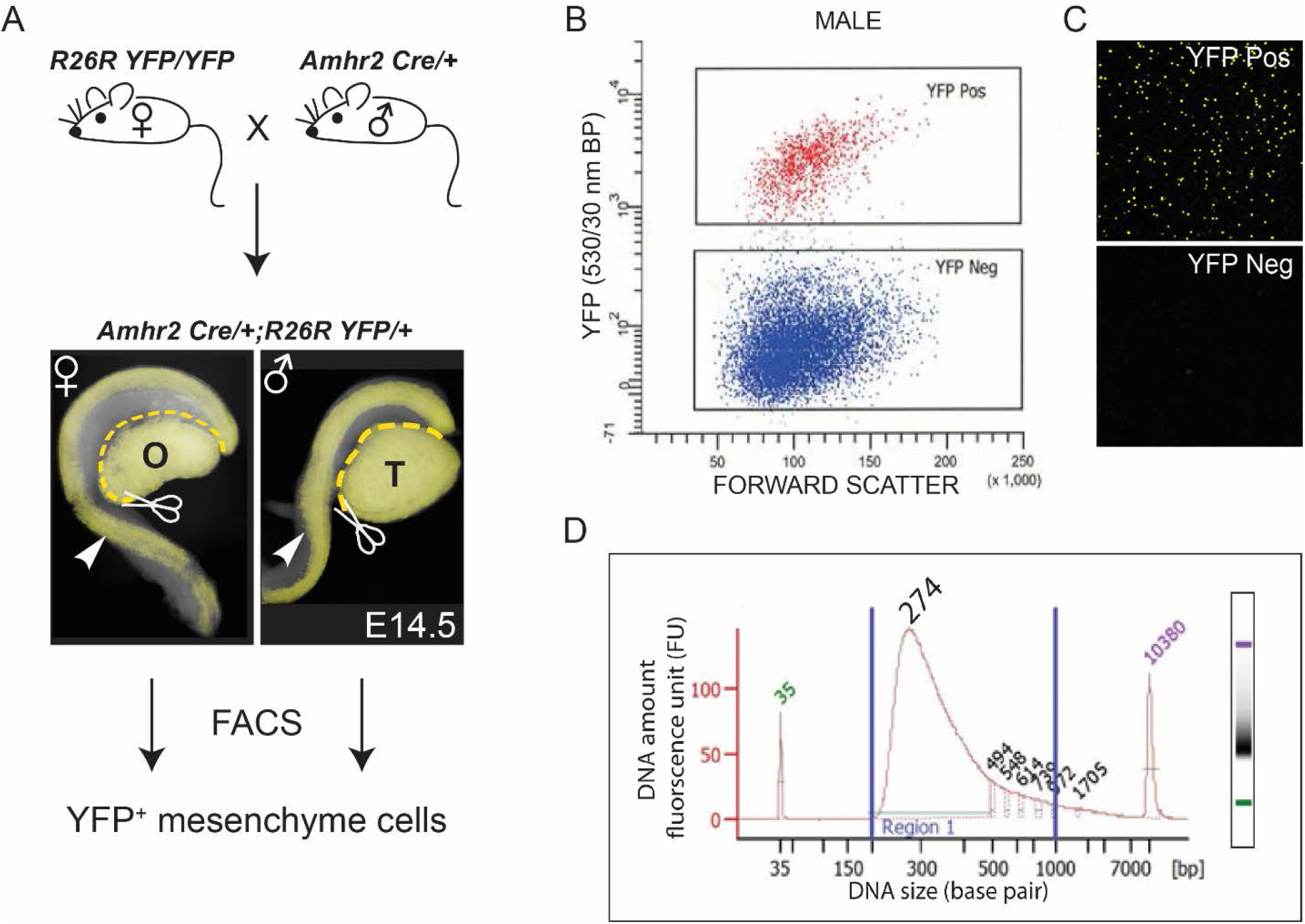
A transcriptome screen for male-specific genes expressed in *Amhr2-expressing* Müllerian duct mesenchyme. **A,** Breeding scheme to isolate *Amhr2*-marked Müllerian duct (MD) mesenchyme cells from E14.5 male and female embryos. The *Amhr2-Cre* allele is expressed in MD mesenchyme cells and somatic cells of the gonad. Mesonephroi containing the MD (arrowheads) and Wollfian ducts were isolated by removing the gonads. Subsequently, YFP^+^ MD mesenchyme cells were isolated by FACS. **B,** Representative plot forward scatter versus YFP emission (530/30 nm band pass filter) from 5 male *Amhr2*^*Cre*/+^; *R26R*^*YFP*/+^ mesenophros pairs to isolate YFP^+^ cells. **C,** Representative confocal image of sorted cell fractions **D,** High sensitivity DNA chip analysis of index pooled cDNA libraries for RNA-Seq. Ovary; O, Testis; T.

**Table 1.**
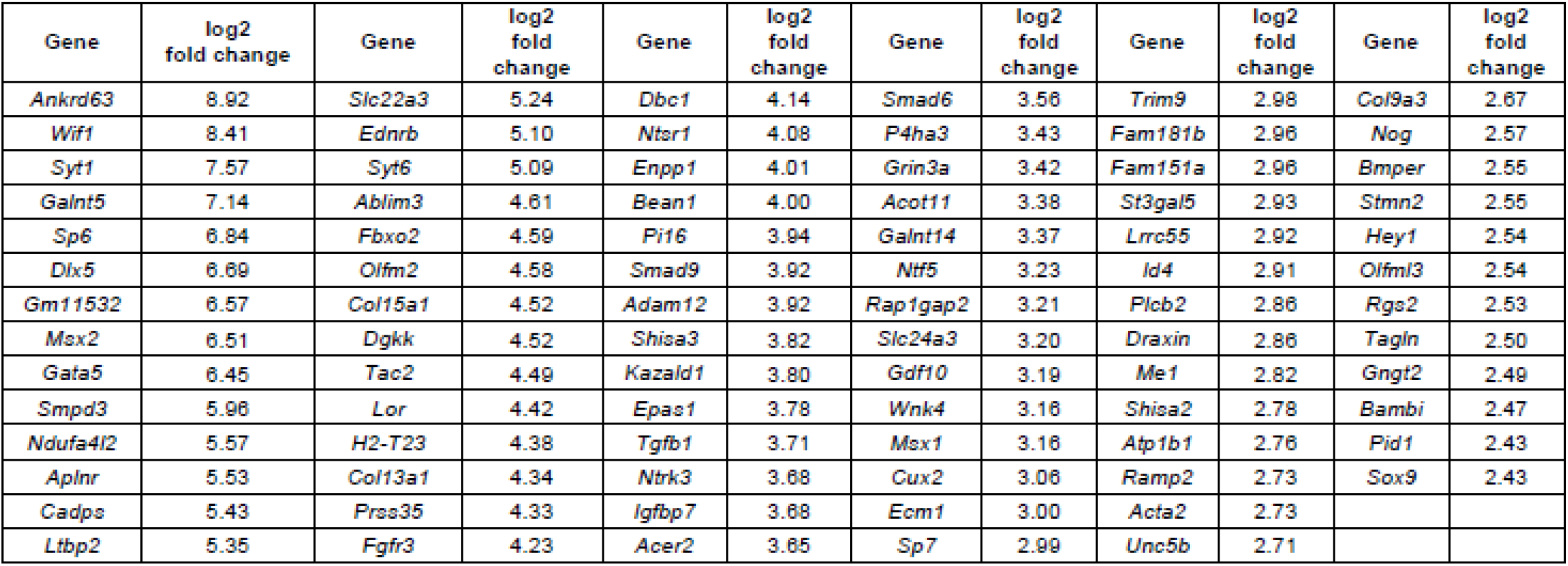
Genes upregulated fn male Müllerian duct mesenchyme.

### *Osterix* is expressed only in male MD mesenchyme during AMH-induced MD regression

OSTERIX (OSX) is a C2H2-type zinc finger transcription factor first cloned in a screen of C2C12 cells induced by BMP2 to differentiate into osteoblasts. *Osx* is required for osteoblast differentiation and bone formation (16). However, to our knowledge this is the first study to identify a role for *Osx* during reproductive tract development. Using *in situ* hybridization we found that *Osx* transcripts are localized specifically in the male MD mesenchyme at E14.5 (**Fig. 2A, B**). To determine whether the OSX protein followed a similar dynamic, we assayed OSX via immunofluorescence and found that OSX is expressed only in male MD mesenchyme at E14.5 indicating a sexually dimorphic expression pattern (**Fig. 2C, D**). This analysis was confirmed by the GUDMAP database (http://www.gudmap.org) wherein *Osx* expression in the MD has a male- specific pattern (17).

**Fig. 2.**
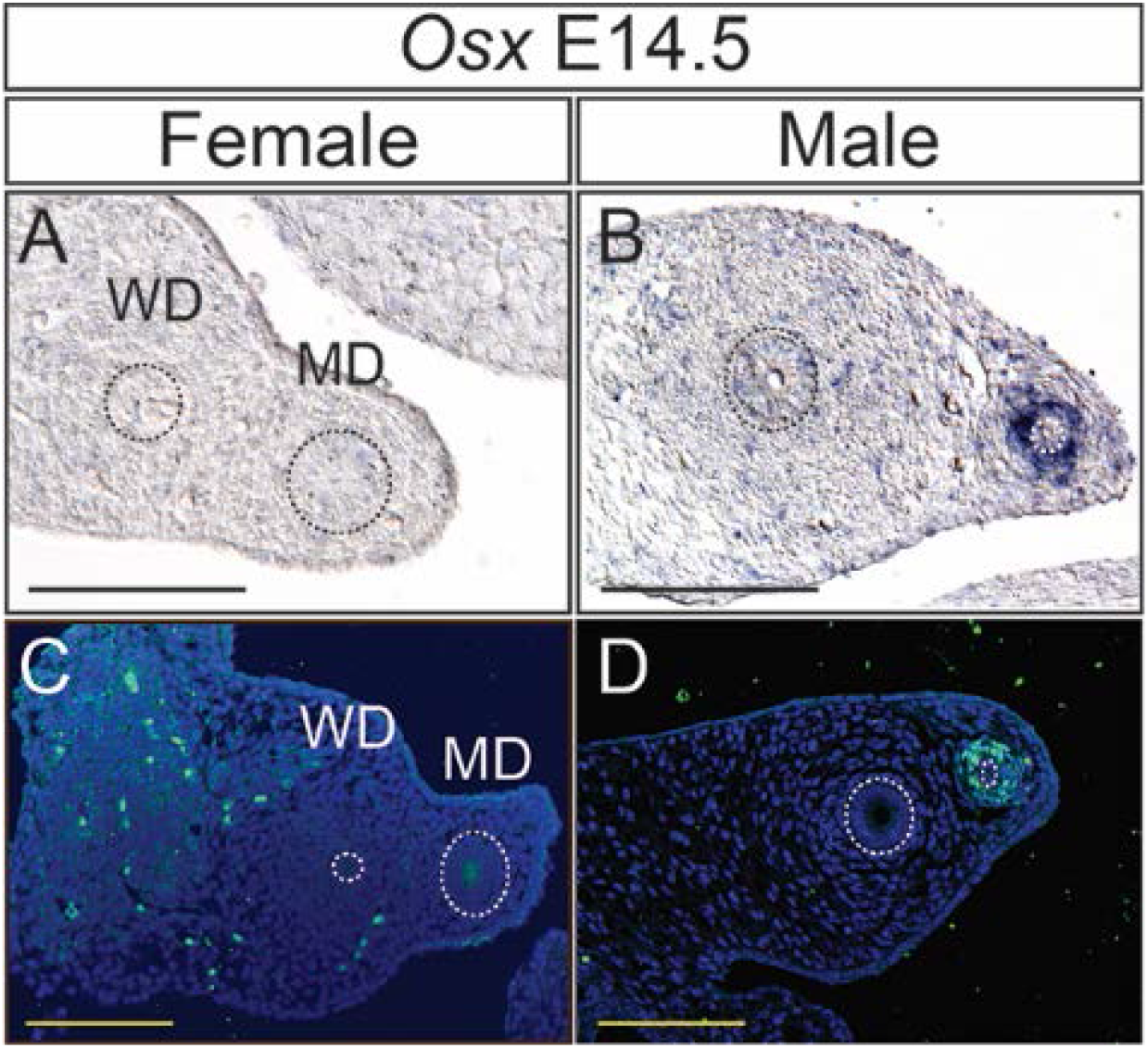
Male-specific expression of Osterix in the mouse Müllerian duct mesenchyme. **A-D,** *In situ* hybridization for *Osx* transcripts (**A, B**) and immunofluorescence for OSX (**C, D**) on paraffin sections of mesonephroi at E14.5. MD and WD epithelium outlined with a dashed line. MD, Müllerian duct; WD, Wolffian duct; scale bar = 100 μm.

Previously, we generated an *Osx-lacZ* (*Z*) null allele in mice that originally demonstrated that *Osx* is required for osteoblast differentiation (16). *Osx-lacZ* expresses beta-galactosidase from the endogenous *Osx* locus and was used to temporally and spatially map expression of *Osx* during MD regression. Whole mount beta-galactosidase staining of E13.5 *Osx*^*Z/+*^ reproductive tracts showed no expression in either males or females (data not shown). Expression of *Osx*^*Z*/+^ begins at around E13.75 in male MD mesenchyme and robust expression throughout the length of the MD is present by E14.5 (**Fig. 3F, G**). This temporal expression pattern is consistent with activation of *Osx* expression by the AMH signaling pathway. While *Amh* transcripts are present in the testis as early as E11.5 in mouse, initiation of AMH signaling in the MD follows expression of *Amhr2* in the MD mesenchyme beginning at ~E13.25 (18, 19). No expression is detected in the female reproductive tract at any time points examined (**Fig. 3A-E**). In males, *Osx*^*Z*/+^ expression is subsequently lost as the MD regresses (**Fig. 3 H-J**). The expression patterns of endogenous OSX and *Osx-lacZ* suggested that *Osx* expression in the MD mesenchyme is mediated by the AMH signaling pathway.

**Fig. 3.**
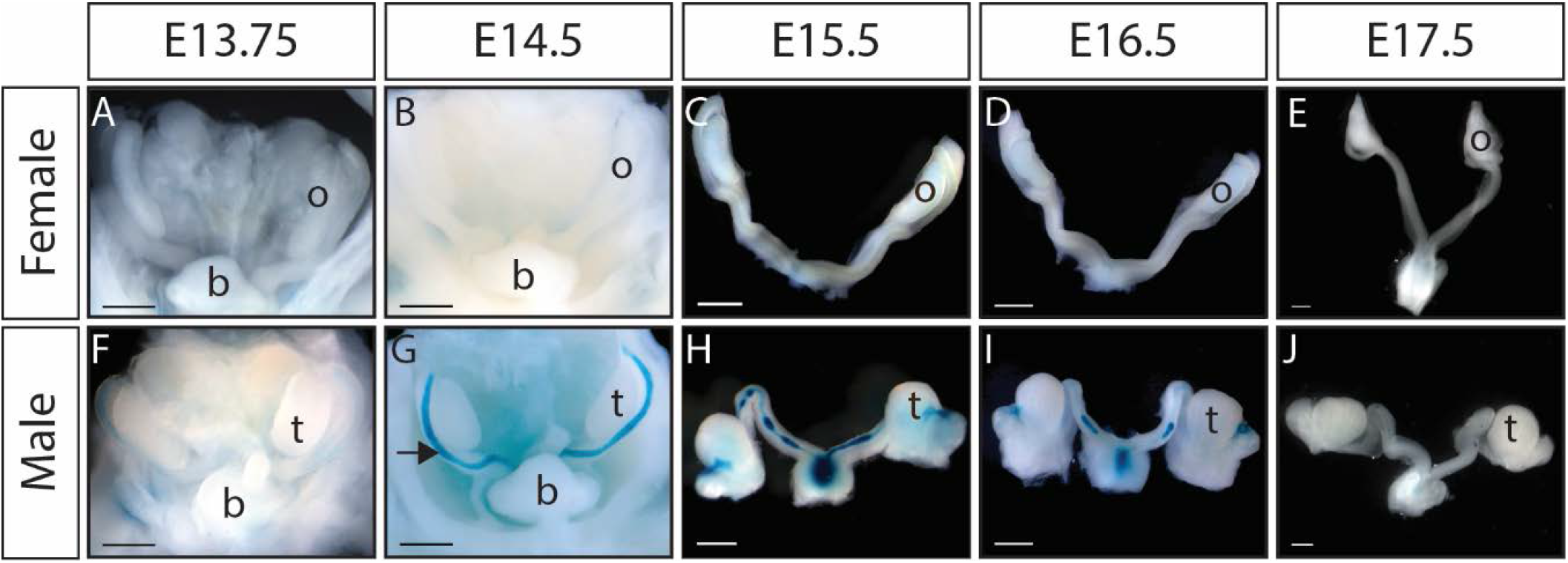
*Osterix-lacZ (Osx-Z)* expression is detectable in Müllerian duct mesenchyme during regression in males but absent in females. **A-J,** Whole mount beta galactosidase staining of *Osx*^*Z*/+^ female and male reproductive tracts. **A-E,** No expression is detected in female embryos from E13.75 to E17.5. **F, G,** *Osx-lacZ* is expressed in male embryos beginning at E13.75 and by E14.5 is expressed in the mesenchyme throughout the length of the MD. **H-J,** As the MD regresses, *Osx-lacZ* expression is also lost. b, bladder; o, ovary; t, testis; scale bar = 500 μm.

### AMH signaling is necessary and sufficient for *OSX/Osterix* expression in MD mesenchyme

To determine if AMH signaling is required for *Osx* expression in the male MD mesenchyme, we examined OSX expression in E14.5 *Amhr2^Z/Cre^* males. *Amhr2^Z/Cre^* is a functional *Amhr2* null genotype and therefore lacks AMH signaling, leading to female reproductive tract development in mutant males (18, 20). While OSX protein can be detected in the MD mesenchyme of wild-type males at E14.5, no detectable OSX is present in *Amhr2^Z/Cre^* males (**Fig. 4A, B**). The ability of AMH signaling alone to activate *Osx* expression was tested using our *MT-hAMH* transgenic mouse model. The *MT-hAMH* mouse broadly expresses human AMH, causing MD regression in females (5). *Osx-lacZ* is not expressed in E14.5 *Osx*^*Z*/+^ females but is induced in the MD mesenchyme of *Osx*^*Z*/+^; *MT-hAMH* females (**Fig. 4C-F**). These results indicate that activation of the AMH signaling pathway is necessary and sufficient to initiate expression of *Osx/OSX* in the MD mesenchyme during reproductive tract differentiation.

**Fig. 4.**
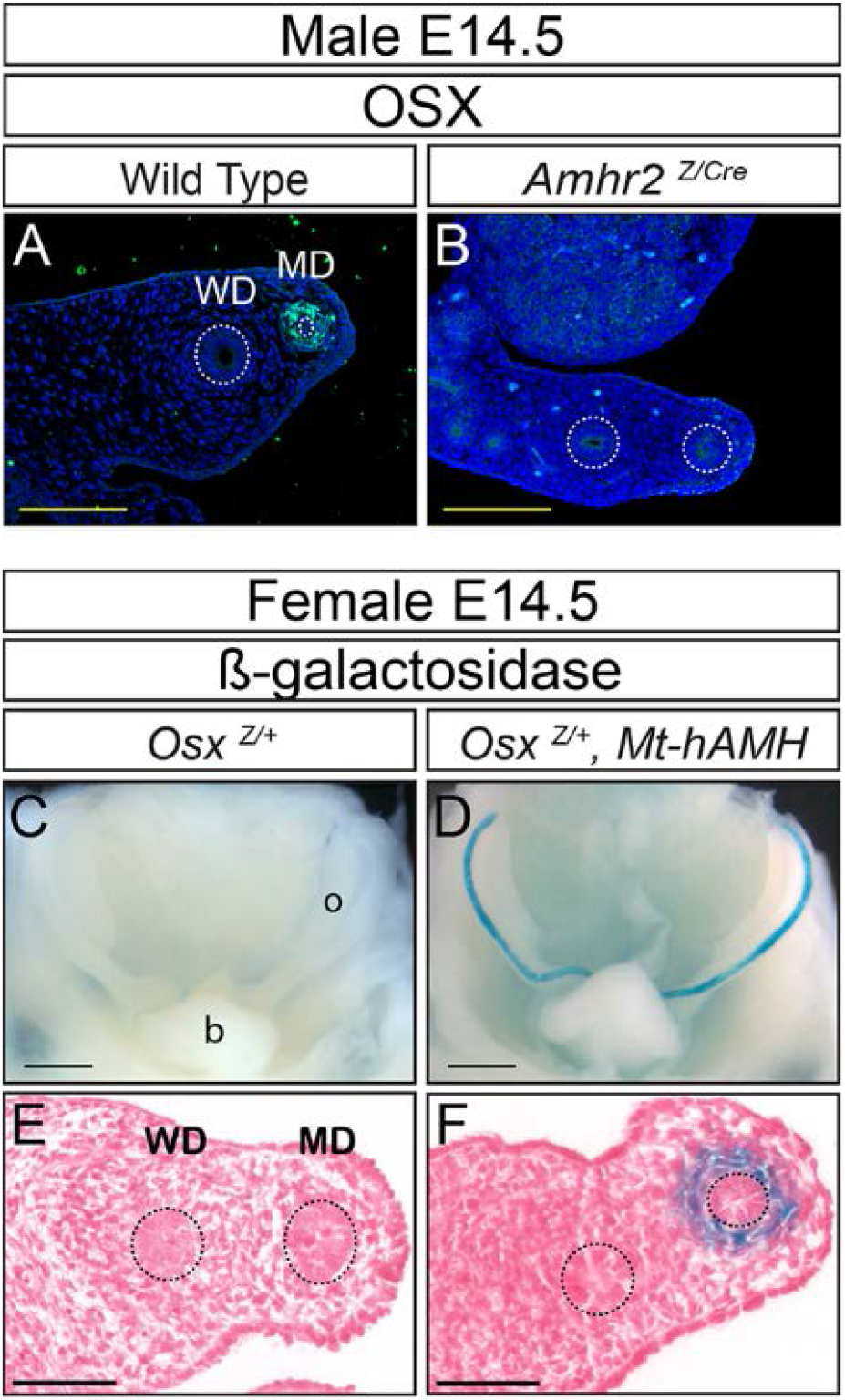
AMH signaling is necessary and sufficient for OSX expression in the MD mesenchyme. **A, B,** Immunofluorescence for OSX on paraffin sections of mesonephroi at E14.5. OSX is not expressed in *Amhr2*^*z/Cre*^ males, indicating AMH signaling is required for OSX expression in the male MD. **C-F,** Beta galactosidase staining of *Osx*^*z*+^ and *Osx*^*Z*/+^; *Mt-hAMH* female reproductive tracts. Activation of the AMH signaling pathway in the female reproductive tract is sufficient to induce expression of *Osx-lacZ* in the MD. MD and WD epithelium outlined with a dashed line. b, bladder; MD, Müllerian duct; o, ovary; WD, Wolffian duct; **A, B, E, F,** scale bar = 100 μm; **C, D,** scale bar = 500 μm.

### Beta-catenin is not required for OSX expression in MD mesenchyme

Beta-catenin functions downstream of AMH signaling and is required to regulate MD regression during male sex differentiation (10). However, the downstream molecular targets of beta-catenin in the MD mesenchyme are unknown. *Osx* expression is upregulated by activated beta-catenin in osteoblasts, cementoblasts, and long bones (21, 22). In addition, the proximal promoter of *Osx* can be activated by beta-catenin in *in vitro* luciferase assays and β-catenin directly binds the first intron of *Osx* in osteoblasts (21, 23). Therefore, we wanted to determine if *Osx* is a downstream target of beta-catenin in the MD mesenchyme and possibly a contributing factor to the complete block in MD regression observed in males with MD mesenchyme knockout of *beta-catenin*. Reproductive tracts of *Amhr2*^*Cre*/+^; *Ctnnb1^flox/flox^* male embryos lack expression of *beta-catenin* in the MD mesenchyme and were examined at E14.5 for OSX expression by immunofluorescence. *Amhr2*^*Cre*/+^; *Ctnnb^flox/flox^* male embryos expressed OSX protein in the MD mesenchyme at E14.5 (**Fig. 5 A, B**). Immunofluorescence of E14.5 male *Osx^Z/Z^* reproductive tracts lacked signal as expected and were used to confirm the specificity of the OSX antibody (data not shown). These results suggest that β-catenin is not required for the expression of OSX in the MD mesenchyme during male sex differentiation. Given that we established *Osx* is a downstream target of AMH signaling but not a direct target of known AMH effector beta-catenin we next analyzed the influence of *Osx* loss on MD regression *in vivo*.

**Fig. 5.**
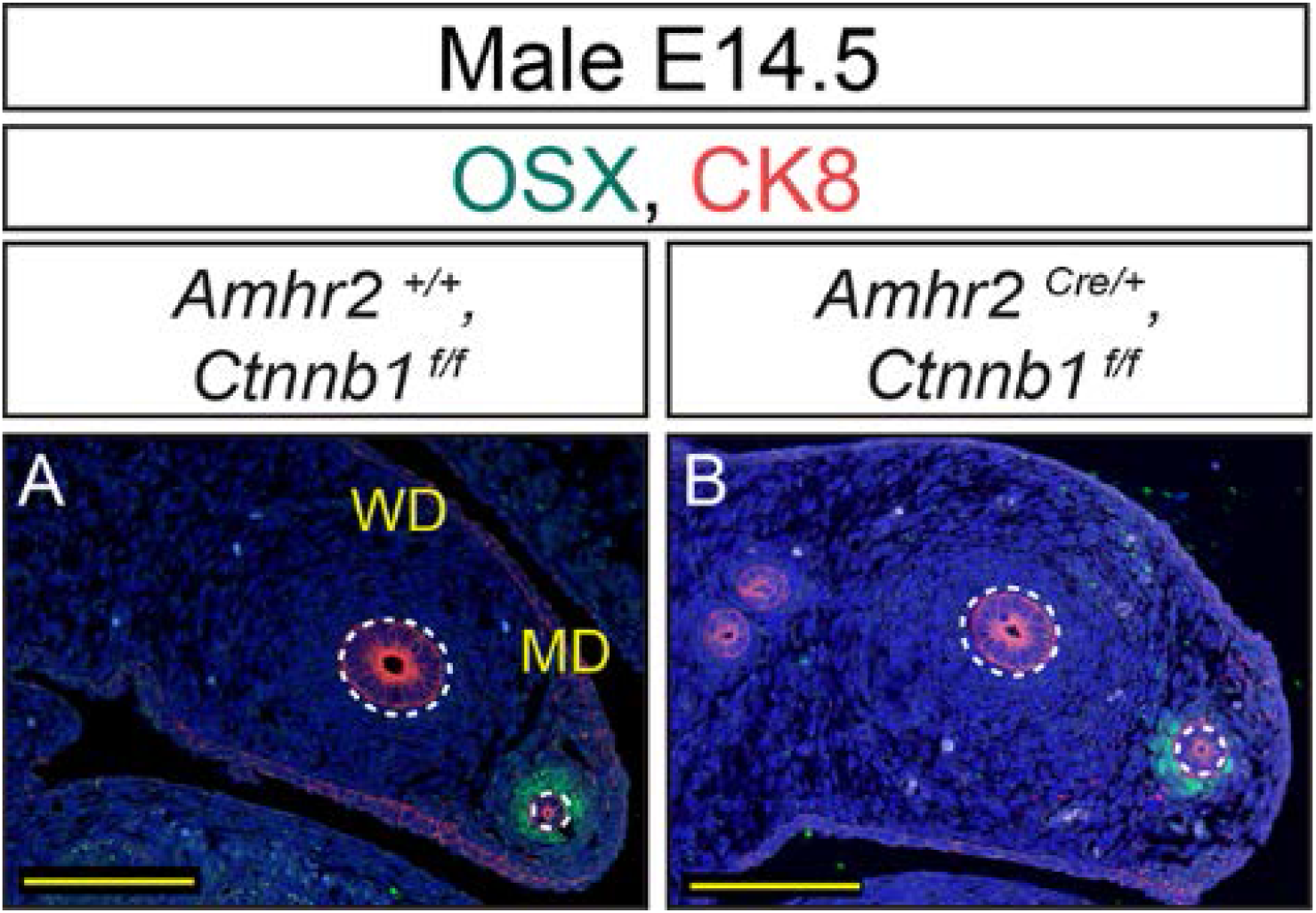
β-catenin is not required for OSX expression in the MD mesenchyme. **A, B,** Immunofluorescence for OSX on paraffin sections of mesonephroi at E14.5. OSX is expressed in *Amhr2*^*Cre*/+^; *Ctnnb1^flox/flox^* males, indicating β-catenin is not required for OSX expression in the male MD. MD and WD epithelium outlined with a dashed line. MD, Müllerian duct; WD, Wolffian duct; scale bar = 100 μm.

### MD regression is delayed in *Osx* null male mice

*Osx* null mice die at birth with severe skeletal defects (16). However, MD regression is complete in males at the time of birth. To determine if *Osx* is required for MD regression we examined *Osx^Z/Z^* embryos at E15.5, E16.5, E17.5 and E18.5. At E15.5, the MD of *Osx^Z+^* and *Osx^Z/Z^* males are intact. However, the *Osx*^*Z*/+^ MD is thinner than the *Osx^Z/Z^* MD, indicating a delay in MD regression in the *Osx^Z/Z^* males (data not shown). At E16.5, the MD of *Osx*^*Z*/+^ males have large gaps. However, the MD of *Osx^Z/Z^* males retain longer lengths of intact MD and fewer gaps are observed (**Fig. 6A, B**). At E17.5, longer MD tissue remnants are observed near the testis in *Osx^Z/Z^* males in comparison to *Osx*^*Z*/+^ males (**Fig. 6C, D**). At E18.5 *Osx^Z/Z^* males have a normal pattern of MD regression and the delay in MD regression defects is resolved. Other developmental delays or growth defects were not evident by gross examination of *Osx* null males (data not shown). This ~24-hour delay in MD regression was observed in all *Osx* null mutant males examined (n=3). This suggests that *Osx* contributes to MD regression. *Osx* has well-established roles in osteoblast, odontoblasts, and cementoblasts differentiation (16, 24). However, this is the first evidence for *Osx* function in sex differentiation.

**Fig. 6.**
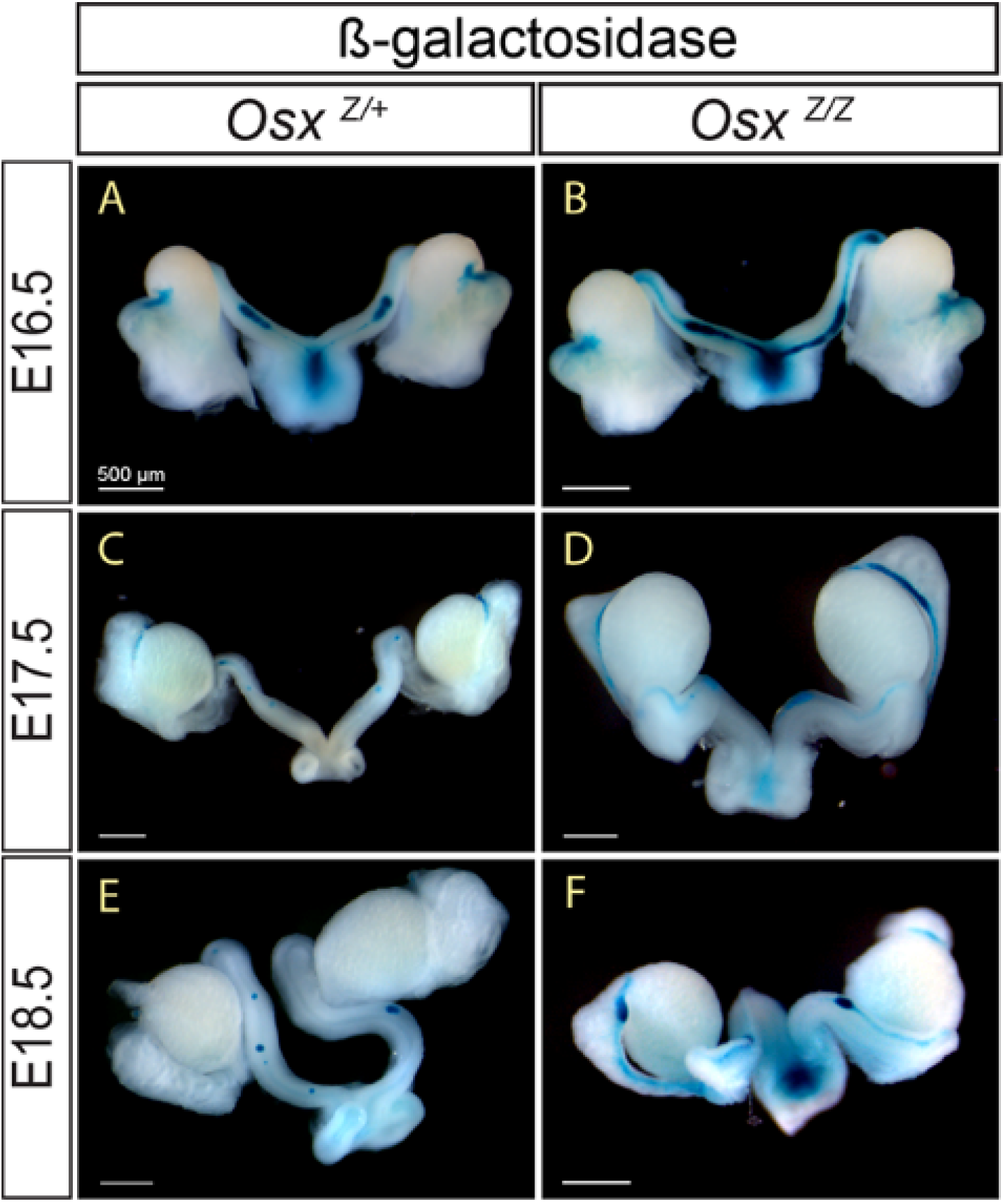
Progression of MD regression is delayed in *OSX^Z/Z^* null mice. **A-F,** Whole mount beta galactosidase staining of *Osx*^*z*/+^ (**A, D, E**) and *Osx^Z/Z^* (**B, D, F**) male reproductive tracts. **A, B,** E16.5 *Osx^Z/Z^* males have longer lengths of MD remnants in comparison to *Osx*^*Z*/+^ males. **C, D,** E17.5 MD tissue remnants are observed near the testis in *Osx^Z/Z^* males. **E, F,** E18.5 *Osx^Z/Z^* regression defects are resolved and appear similar to *Osx*^*Z*/+^ males. Scale bar = 500 μm. Three or more of each genotype were observed for each developmental time point.

## Discussion

We report on a novel AMH-induced gene, *Osx*, as a regulator of MD regression identified using a global unbiased approach. The identification of *Osx* in this process adds a key link to the gene expression network that may underlie MD regression. During development in skeletal bone and tooth, *Osx* has been demonstrated to be a key mesenchymal factor necessary for cell-fate decisions in the differentiation of specialized cells (16, 25). To our knowledge ours is the first study implicating *Osx* as a factor in sex differentiation.

*Osx* is an important factor in bone homeostasis in adults and has roles in degradation of cartilage matrix by directly transcriptionally activing *matrix metalloproteinase, Mmp13* (26). Further, in metastatic breast cancer, modulation of *Osx* levels results in a corresponding change in MMP-2 levels (27). Two important processes in MD regression are the breakdown of the basement membrane and apoptosis of MD epithelial cells. Secreted MMPs function to degrade extracellular matrix and also cleave signaling molecules to control multiple cellular processes including differentiation and apoptosis (28). MMP-2 has been identified as a downstream target of AMH signaling and is expressed in a male-specific pattern in the MD during regression. Knockdown of *Mmp2* using a morpholino and MMP inhibitors block MD regression in *in vitro* organ culture experiments (29). However *Mmp2* null male mice regress the MD, suggesting other MMPs may compensate for loss of MMP-2 (29). MMP-14 is expressed in the reproductive tract during MD regression and may compensate for MMP-2 but its expression is not sexually dimorphically (29). Interestingly a female patient that lacks a uterus has been reported with a duplication in chromosome 14 containing the *MMP14* gene region (30). Considering that *Osx* modulates MMP expression in other systems, *Osx* may be contributing to MD regression by regulating MMP levels.

The AMH signaling pathway shares type one receptors, ACVRI and BMPR1A, and downstream SMADs with the BMP pathway. This suggests that AMH-induced expression of *Osx* in the reproductive tract may be regulated by similar mechanisms as its regulation by BMP signaling in bone and tooth differentiation. During skeletal and tooth root formation, BMP signaling is capable of inducing *Osx* expression by both *Runx2*-dependent and -independent mechanisms. In our study, transcript levels of *Runx2* are not differentially expressed in the MD mesenchyme at E14.5 between males and females, suggesting in male reproductive tract differentiation *Osx* expression is independent of *Runx2*. It is possible *Runx2* is expressed in the male reproductive tract prior to activation of *Osx* expression and later downregulated by E14.5 or that male-specific co-factors of RUNX2 are required to induce *Osx* expression. In addition to *Runx2, Osx* expression is upregulated by activated BETA-CATENIN in osteoblasts, cementoblasts, and long bones (21, 22). In our previous study we found that *β-catenin* in the MD mesenchyme is required for AMH-induced regression of the MD. However our results show OSX expression in the MD mesenchyme does not require *β-catenin*. Because *β-catenin* conditional knockout mice have complete retention of the MD and *Osx* null males only show a delay in MD regression it is unlikely OSX is required for *β-catenin* expression in the MD mesenchyme. Of note the chromosomal location of the *Osx* gene is adjacent to the *Amhr2* gene and this linkage is conserved in mammals. This implies the potential for shared enhancer/regulatory elements for MD mesenchyme expression.

Interestingly, Park et al. (2014) performed a related study but used microarrays to determine transcript profiles of whole E14.5 mesonephroi from wild-type and *Amhr2* mutant males to identify AMH-induced genes. Our study (MD mesenchymal cells only) and the Park study (whole mesonephroi) resulted in very similar lists of genes up-regulated in male MD (12). This suggests gene activation that mediates MD regression is primarily occurring in the MD mesenchyme.

Loss of OSX protein results in a ~ 24-hour delay in the progression of MD regression. The spatial pattern of MD regression in *Osx* null male embryos was comparable to what we observed in heterozygous littermates with gaps apparent throughout the MD by E16.5. However, the length of remaining MD was increased in the absence of *Osx* at E16.5 & E17.5. By E18.5 the delay in MD regression was resolved. This suggests that OSX may play an important role in the timing of initiation of MD regression. It is possible OSX is an early activator of molecular signals necessary for clearance of MD cells by apoptosis or other mechanisms during regression. After a brief delay due to the loss of initial activation by OSX, these clearance signals are then activated by other as of yet unidentified AMH-induced factors.

It is also possible that SP family member *Epfn/Sp6* is compensating for the loss of *Osx* during MD regression and allowing MD regression to proceed. There is high expression and log2 fold change of *Epfn* in male MD mesenchyme in our dataset. In addition, there is high homology of EPFN with OSX protein (46% identity match overall/ >90% identity match in DNA binding zinc finger domain) and sequence-based hierarchical clustering place *Epfn* and *Osx* within the same subfamily of SP proteins (31). However, to our knowledge there is no previous evidence for overlapping function. Both *Epfn* and *Osx* have roles in tooth development but expression patterns are not overlapping with *Osx* expressed in dental mesenchyme and *Epfn* expressed in dental epithelial cells (25). In the MD mesenchyme, both *Epfn* and *Osx* are expressed in males.

The MD regression defects described have included complete and partial retention of MD derived tissues in postnatal and adult males and delayed regression that resolves at later embryonic stages. Factors whose loss results in no regression of the MD include *Wnt7a* (required for *Amhr2* expression), *Amh, Amhr2, Bmpr1a/Acvr2, Smad1,5,8*, and *β-catenin*. With the exception of *β-catenin* all of these genes function in the initial activation of the AMH signaling pathway (2). Partial retention of MD-derived tissues has been associated with reductions in levels of AMH hormone and AMHR2, leading to AMH signaling below a threshold level needed for complete MD regression (32, 33). Similarly, mice with lags in testis development and consequent delays in AMH secretion also exhibit deferred MD regression (34). Partial retention of MD remnants was also observed in conditional knockout (CKO) in the MD mesenchyme of the TGF-β signaling pathway common Smad (co-Smad) *Smad4*. The defect in MD regression in the CKO *Smad4* males is hypothesized to be caused by an observed reduction in *β-catenin* (35). Interestingly constitutive activation of *β-catenin* in the MD mesenchyme resulted in a similar partial retention of MD tissues in adult males (36). To our knowledge the delay in MD regression in the *Osx* null males is the first described that is independent of AMH signaling levels and *β-catenin* expression.

Numerous studies have examined the consequences of gene mutations on MD regression in postnatal or adult males and considered any mutants without retained uterine tissue to have “normal” MD regression. This approach works well to identify factors required for activation of AMH signaling during MD regression such as *Wnt7a, Amh* and *Amhr2* or early downstream effectors in the signaling cascade such as SMADs 1, 5, 8 (2). However, monitoring mutant males for retained MD-derived tissues after MD regression is complete does not identify potential changes in the timing or process of MD regression. By examining *Osx* null mutant males at early time points during MD regression, we were able to identify a potential role for *Osx* in the initiation of MD regression that would have been missed if we had only examined newborn males. Considering the results of this study, it may be of benefit to revisit candidate factors without postnatally retained uterine tissue at earlier time points during MD regression.

## Materials and Methods

### Mice

*Ctnnb1^flox/flox^* mice were obtained from the Jackson Laboratory (Bar Harbor, ME, USA) and maintained on a C57BL/6J × 129X1/SvJ × 129S1/Sv mixed background (37). *Amhr2^tm3(Cre)Bhr^* (*Amhr2-Cre*; (20)), *Gt(ROSA)26Sor^tm1(EYFP)Cos^* (*R26R-YFP*; (38)), and *Amhr2^tm2Bhr^* (*Amhr2-lacZ*; (18)) were maintained on a C57BL/6J × 129/SvEv mixed background. *Sp7^tm1Crm^* (*Sp7-lacZ*; (16)), and *Mt-hAMH*^*tg*/+^ (*Mt-hAMH*; (7)) were maintained on a C57BL/6J congenic background. *Amhr2-Cre, R26R-YFP, Amhr2-lacZ, Sp7-lacZ, Ctnnb1^flox/flox^*, and *Mt-hAMH*^*tg*/+^ mice were genotyped as previously described (7, 8, 16, 18, 37, 38). The Institutional Animal Care and Use Committee of the University of Texas MD Anderson Cancer Center approved all animal procedures. Experiments were performed in agreement with the principles and procedures outlined in the National Institutes of Health Guidelines for the Care and Use of Experimental Animals.

### Isolation of MD mesenchyme RNA

E14.5 *Amhr2*^*Cre*/+^; *R26R*^*yfp*/+^ mesonephroi were digested in 250 *μ*L 0.25% Trypsin-EDTA (Gibco# 25200-056) for 20 minutes and mechanically dissociated into single cell suspensions and filtered through 35 μm Falcon Tubes with Cell Strainer Cap (Fisher Scientific, US). YFP+ Müllerian duct mesenchymal cells were directly sorted into TRI reagent RT-LS (Molecular Research Center Inc., Cincinnati, OH, USA) using a BD FACS Aria I cell sorter (BD Biosciences, USA) and homogenized. Total RNA was isolated following the manufacturer’s recommended protocol with the following modifications; PelletPaint (EMD Millipore, Billerica, MA, USA) was added to samples to visualize RNA pellets and an additional ethanol precipitation was performed to remove any remaining phenol. RNA integrity was measured with an Agilent 2100 Bioanalyzer (Agilent Technologies, Santa Clara, CA, USA), and a sample with an RNA Integrity Number greater than or equal to 8 was considered acceptable.

### Generation of RNA-Seq cDNA libraries

cDNA libraries were generated using the TruSeq v2 kit (Illumina, San Diego, CA,USA) following the manufacturer’s recommended protocol from three biological replicates each of male and female. Each replicate consisted of ≥ 100 ng of total RNA isolated from pooled YFP+ Müllerian duct mesenchyme cells from 2-7 embryos. Individual library quality was assessed by high sensitivity DNA Bioanalyzer chip (Aligent, Santa Clara, CA) and the concentration of each library was determined using the Qubit picogreen assay (Life Technologies, Grand Island, NY). Equal quantities of each indexed library were pooled and sequenced on the Illumina HiSeq 2000 platform to generate paired end 76 base pair reads. Raw data files for the RNA sequencing analysis have been deposited in the NCBI Gene Expression Omnibus (GEO) with the accession number GEO: XXXXXXX.

### Bioinformatic analysis of transcriptomes

RNA-seq data was mapped to the mouse genome with the bioinformatics tools Tophat and Bowtie using reference genome build NCBIM37 (13). Ensembl annotation for the mouse genome was downloaded from the iGenome website (http://support.illumina.com/sequencing/sequencing_software/igenome.ilmn). Mapped RNA-seq data were then subject to differential expression analysis. Two bioinformatics tools were used for the analysis: Cufflink (13) and DESeq (14). Cufflink uses mapped reads to generate a parsimonious set of transcripts, and then estimates the abundance of the transcripts (39). DESeq utilizes the raw counts of mapped reads as data input and uses the negative binomial distribution model (14). Using both methods, a list of genes with significant differential expression was obtained. To study the possible functions related to the changes of gene expression, pathway analysis was performed using Ingenuity Pathway Analysis tool from Ingenuity Systems (Ingenuity^®^ Systems, www.ingenuity.com).

### In situ hybridization

Embryos and reproductive tissues from neonates or adults were processed and section *in situ* hybridization was performed as previously described (10). *Osx* RNA probe was generated from plasmid pBSOsxBP containing a 560 bp fragment of mouse *Osx* (*Osx* transcript NM_1330458.4 bp 106-672) (16).

### X-gal staining

Embryos were collected at stages E13.5 to E18.5. Lower body trunks with urogenital tissues were isolated and processed for X-gal staining to visualize lacZ expression as described (18).

### Immunofluorescence

The following primary antibodies were used for immunofluorescence: Cytokeratin 8 (1:100; Developmental Studies Hybridoma Bank; TROMA-I) and OSTERIX (1:50; a gift from Benoit de Crombrugghe) (16). Immunofluorescence was performed as previously described (40). At least 3 specimens of each genotype were analyzed for each staining.

## Acknowledgements

We thank Swathi Arur, Zer Vue, Shuo-Ting Yen, and Alejandro Elder Ontiveryos for helpful discussions. pBSOsxBP and OSX antibody were kindly provided by Benoit de Crombrugghe. Supported by National Institutes of Health (NIH) grant HD030284 and the Ben F. Love Endowment to R.R.B. R.D.M. was supported by NIH T32 grant CA009299 and a postdoctoral fellowship from UTMDACC Center for Stem Cell and Developmental Biology. Veterinary resources, flow cytometry, and DNA sequencing were supported by NIH grant CA16672.

